# A Cross-Modal Autoencoder Framework Learns Holistic Representations of Cardiovascular State

**DOI:** 10.1101/2022.05.26.493497

**Authors:** Adityanarayanan Radhakrishnan, Sam Freesun Friedman, Shaan Khurshid, Kenney Ng, Puneet Batra, Steven Lubitz, Anthony Philippakis, Caroline Uhler

## Abstract

A fundamental challenge in diagnostics is integrating multiple modalities to develop a joint characterization of physiological state. Using the heart as a model system, we develop a cross-modal autoencoder framework for integrating distinct data modalities and constructing a holistic representation of cardiovascular state. In particular, we use our framework to construct such cross-modal representations from cardiac magnetic resonance images (MRIs), containing structural information, and electrocardiograms (ECGs), containing myoelectric information. We leverage the learned cross-modal representation to (1) improve phenotype prediction from a single, accessible phenotype such as ECGs; (2) enable imputation of hard-to-acquire cardiac MRIs from easy-to-acquire ECGs; and (3) develop a framework for performing genome-wide association studies in an unsupervised manner. Our results provide a framework for integrating distinct diagnostic modalities into a common representation that better characterizes physiologic state.

## 1 Introduction

Clinicians leverage measurements across many complementary diagnostic modalities to develop an integrated understanding of a patient’s physiological state. For example, heart function can be interrogated with a variety of modalities, such as electrocardiograms (ECGs) that provide myoelectric information (e.g. sinus rhythm, ventricular rate, etc.), and cardiac magnetic resonance images (MRIs) that provide structural information (e.g. left ventricular mass, right ventricular end-diastolic volume, etc.). By utilizing measurements across both modalities, we can gain a more holistic view of cardiovascular state than with either modality alone. The recent availability of large-scale cross-modal patient measurements in biobanks [1,2] provides the opportunity to develop systematic and rich representations of physiology. Using the heart as a model system, we here develop such an integrative framework and show its effectiveness in downstream tasks including phenotype prediction, modality translation, and genetic discovery.

Our approach relies on a class of machine learning models called autoencoders. Autoencoders [3, 4] are a class of generative models that serve as a standard method for learning representations from unlabelled data. These models have been successfully applied in a variety of applications including computer vision [5–7], chemistry [8], and biology [9–14]. A line of recent works utilize autoencoders to learn joint representations of multi-modal data including natural images and captions in computer vision [7, 15– 9], nuclear images and gene expression in biology [13], and paired clinical measurements [20–22]. Indeed, autoencoders have been observed to perform competitively with other multi-modal integration methods including classical integration approaches using canonical correlation analysis [23–25] and generative adversarial networks [26, 27]. Unlike these prior works that focus primarily on integrating images and vectorized data such as gene expression, we aim to integrate complex modalities with a temporal element (cardiac MRI videos and ECGs). In addition, we aim to learn a cross-modal representation that can also be used for characterizing genotype-phenotype associations. While various prior works have conducted genome-wide association studies (GWAS) to identify single nucleotide polymorphisms (SNPs) associated with cardiovascular diseases [28, 29], features measured on ECGs [30, 31], or features measured on cardiac MRI [32, 33], these GWAS approaches have relied on labelled data derived from individual modalities and thus the resulting genetic associations are modality specific, i.e., SNPs affecting an ECG would not necessarily be significant on a GWAS for an MRI derived phenotype.

Utilizing cardiac MRI and ECG samples from the UK Biobank [1], we develop a cross-modal autoen-coder framework for building a representation of patient cardiovascular state (Fig. 1a). We show that these learned representations improve phenotype prediction (Fig. 1b). Additionally, our cross-modal autoencoders enable generating hard-to-acquire MRIs from easy-to-acquire ECG samples, and we show that these generated MRIs capture common MRI phenotypes (Fig. 1c). We show that a GWAS on phenotype labels derived from cross-modal embeddings leads to the recovery of known genotype-phenotype associations. Importantly, our framework also allows to perform GWAS in the absence of labeled phenotypes to identify SNPs that generally impact the cardiovascular system. (Fig. 1d).

**Figure 1:**
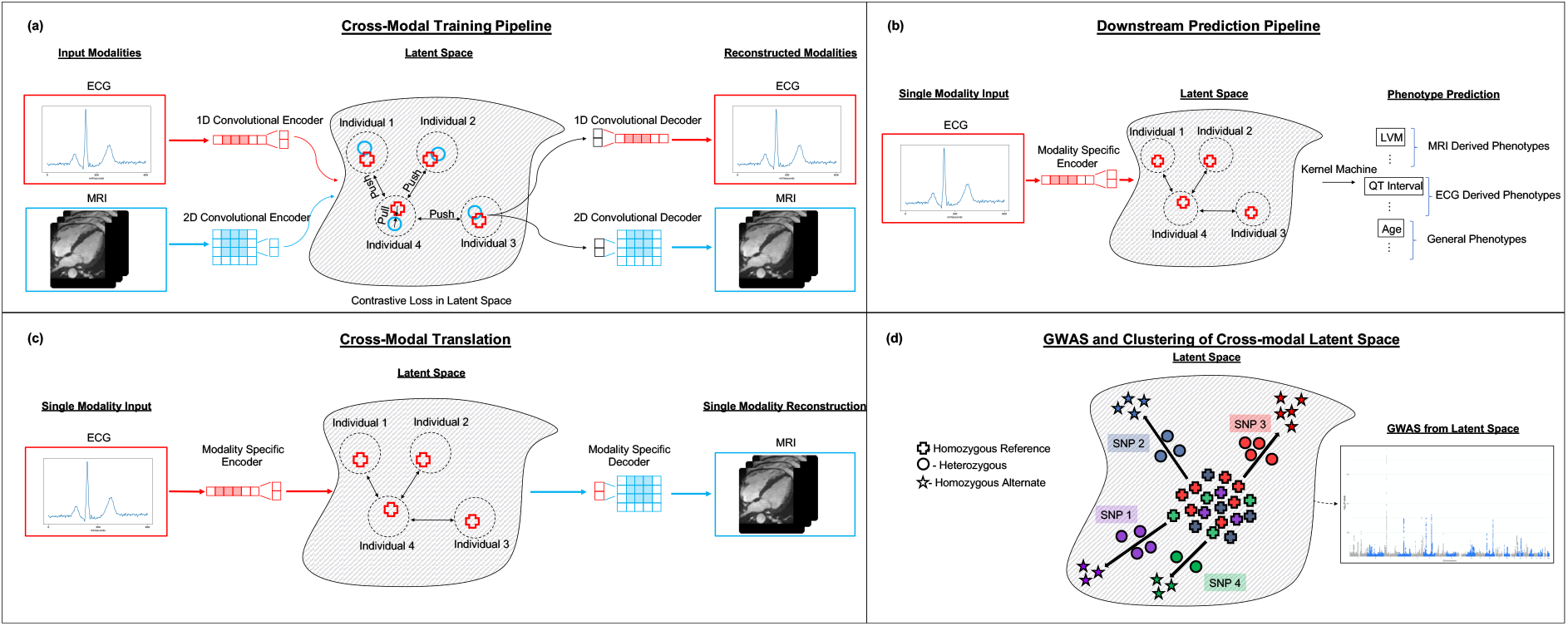
An overview of our cross-modal autoencoder framework for integrating cardiovascular data modalities. Our model is trained on ECG and cardiac MRI pairs from the UK Biobank. (a) A visualization of our training pipeline. Modality specific encoders map data modalities into a shared latent space in which a contrastive loss is used to enforce the constraint that paired samples are embedded nearby and further apart from other samples. Modality specific decoders are then used to reconstruct modalities from points in the latent space. (b) Learned cross-modal representations are used for downstream phenotype prediction tasks by training a supervised learning model (e.g., a kernel machine) on the latent representations. (c) Our framework enables translation between modalities: ECGs can be translated to corresponding MRIs and vice-versa. (d) The learned cross-modal representations can be used to understand genotype-phenotype maps in the absence of labeled phenotypes by performing a GWAS in the cross-model latent space and clustering SNPs via their signatures (i.e., the direction from homozygous reference to the mean of heterozygous and homozygous alternate); SNPs 1 and 4 have similar signatures in the latent space and thus similar phenotypic effects.

## Results

### Cross-modal Autoencoder Framework Enables the Integration of Cardiovascular Data Modalities

To build a cross-modal representation of patient cardiovascular state, we utilize autoencoders to map paired cardiovascular data modalities, i.e. 38,686 paired median 12-lead ECGs and 50 frame videos of long axis cardiac MRIs, from the UK Biobank [1] into a common latent space. A description of the data used in this work is provided in Data Availability 1.5. Building on the traditional autoencoder framework, we train modality-specific encoders and decoders to map to and from this latent space such that the reconstructed training examples are similar to the original examples for all modalities (see Fig. 1a). Additionally, given an ECG and MRI pair for a single individual, we utilize a loss function that ensures that paired ECG and MRI samples are represented via nearby points in the latent space (i.e., using a *contrastive loss*). Importantly, while our model is trained on paired modalities, the model can be applied in settings where only one modality is present. Namely, we simply utilize the embedding given by the trained encoder for the single input modality. A description of our loss function, architectures, and training procedures is given in Methods 1.2 and Supplementary Fig. S1. As indicated in Fig. 1, the resulting representations are useful for a variety of downstream tasks including phenotype prediction, modality translation, and genetic discovery.

### Cross-modal Representations Enable Improved Phenotype Prediction

We first demonstrate that supervised learning on cross-modal representations improves performance on phenotype prediction tasks. While our model is trained on ECG and MRI pairs, we consider the practically relevant setting in which only one modality (e.g. ECG) is available. In this case, we perform supervised learning on embeddings given by a single modality-specific encoder (Fig. 1b). For our cross-modal autoencoder trained on paired cardiac MRI and ECG samples, we show that utilizing standard regression methods (e.g. kernel, linear, or logistic regression) for supervised learning on our cross-modal representations leads to improved prediction of: (1) MRI derived phenotypes (e.g. left ventricular mass, right ventricular end-diastolic volume, etc.) from ECG only; (2) ECG derived phenotypes (e.g. length of PR interval, QT interval, etc.), from MRI only; and (3) prediction of general phenotypes (e.g. age, sex, body mass index, etc.) from either ECG or MRI. We observe that predictive models applied to our cross-modal representations generally outperform supervised deep learning models and supervised learning on traditional unimodal autoencoder representations.

#### Cross-modal embeddings allow for matching cardiac MRI and ECG test samples

We begin by verifying that our training methodology provides a latent space in which corresponding ECG and MRI pairs are nearby. Hence, even in the absence of one of the modalities, the cross-modal autoen-coder provides a representation that is characteristic of all available modalities. In Fig. 2a, we provide a t-distributed stochastic neighbor embedding (t-SNE) comparing the unimodal and cross-modal autoencoder latent space representations for 500 paired ECG and MRI test samples. The t-SNE plots demonstrate that the ECG and MRI samples are well-mixing in the cross-modal latent space, while the two are clearly separated in the corresponding unimodal latent space. To quantify the benefit of cross-modal representations, we compute the accuracy that the correct MRI pair lies within the top *k* nearest neighbors (under cosine similarity) for 4752 test ECGs across embeddings from cross-modal autoencoders, unimodal autoencoders, and a baseline where ECGs and MRIs are randomly paired. Fig. 2b demonstrates that cross-modal representations outperform unimodal representations in this task, with the latter performing similarly to the random baseline.

**Figure 2:**
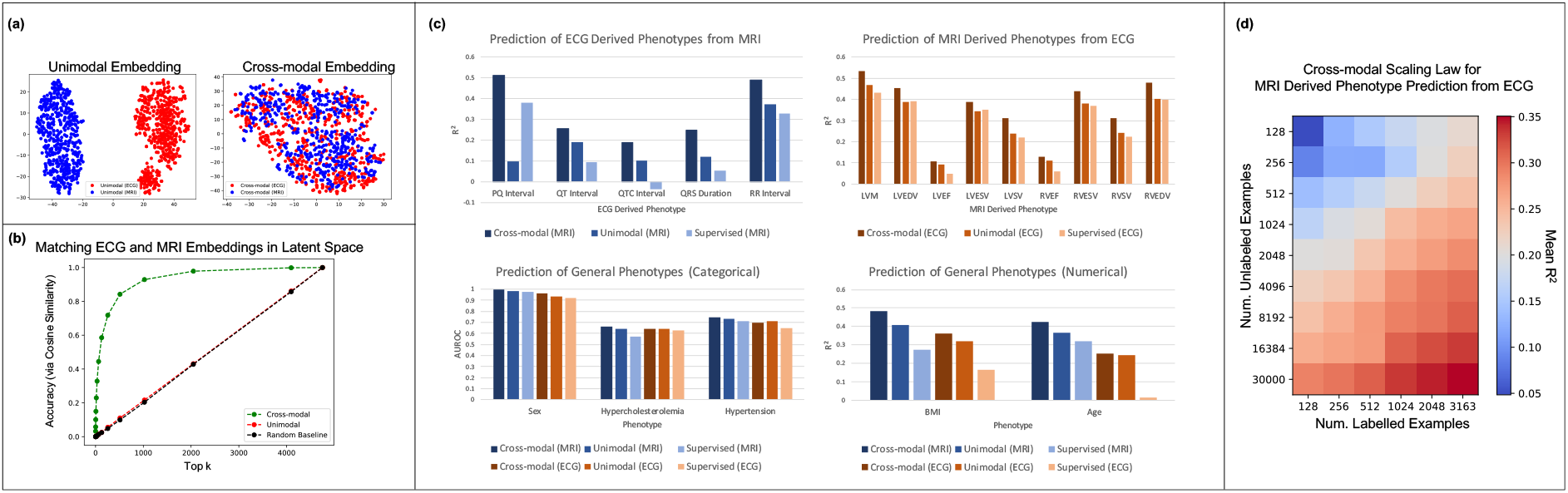
Comparison of the effectiveness of phenotype prediction from cross-modal representations as opposed to unimodal representations or supervised learning from the original modalities. (a) A t-SNE visualization of the cross-modal embeddings for the ECG and MRI samples demonstrates that the modality specifc embeddings are well-mixed, unlike the modality specific embeddings obtained from the unimodal autoencoders. (b) Ranking each MRI by its cosine similarity with a given ECG in the latent space, we visualize the accuracy that the ground truth MRI appears in the top *k* neighbors among 4752 test ECG-MRI pairs from the UK Biobank. (c) Kernel regression on cross-modal representations outperforms kernel regression on unimodal representations and supervised deep learning methods on 4 different tasks: (1) prediction of ECG derived phenotypes from MRIs only; (2) prediction of MRI derived phenotypes from ECG only; (3) prediction of general physiological phenotypes that are of categorical nature from either ECG or MRI; and (4) prediction of general physiological phenotypes that are of continuous nature from either ECG or MRI. (d) Analysis of the scaling law when utilizing our framework for predicting MRI derived phenotypes from ECGs only. We observe that increasing the number of unlabelled ECG-MRI pairs for pre-training boosts the mean *R*^2^ prediction of 9 MRI derived phenotypes by twice as much as increasing the number of labelled MRI samples. This analysis highlights the benefit of collecting more unlabelled ECG-MRI pairs as compared to paired labelled examples for this task.

#### Cross-modal representations improve phenotype prediction from a single modality

We now show that our learned cross-modal representations are more effective for downstream phenotype prediction than unimodal representations or supervised deep learning methods. In particular, we consider 4 groups of phenotype prediction tasks: prediction of (1) continuous valued MRI-derived phenotypes from ECG; (2) continuous valued ECG-derived phenotypes from MRI; (3) categorical physiological phenotypes from either ECG or MRI; and (4) continuous physiological phenotypes from either ECG or MRI (see Fig.2c. For all prediction tasks, we utilized the same training, validation, and held out test data from the UK Biobank. Importantly, we note that all data considered for the downstream prediction tasks were excluded from the training procedure for the cross-modal autoencoders. This is critical since otherwise, we could simply train a cross-modal autoencoder to zero error on paired data, and our learned representations would naturally benefit from using both MRI and ECG features for any downstream prediction task; see Methods 1.3 for details on the data splits considered. Again, we note that only a single modality is used for each of these tasks, i.e., we are not giving the cross-modal autoencoders access to any paired samples for the downstream phenotype prediction tasks.

We utilize kernel regression to perform supervised learning from the cross-modal and unimodal embeddings; see Supplementary Fig. S2 for a comparison with the performance of linear regression and logistic regression. For fair comparison with supervised deep learning models, we extract the embeddings given by the last layer of the trained neural networks and apply kernel regression on these embeddings; see Methods 1.3 for a description of the architectures for all deep networks used in this task. In all but one setting (hypertension classification), we observe that predictions from our cross-modal latent space improve over predictions from unimodal latent spaces and those from direct supervised learning methods. An important practical implication of these results is that our method is capable of improving the prediction of a variety of phenotypes just using ECGs, which are far easier to obtain and more plentiful than MRIs. This is exemplified by the improvement in prediction of MRI derived phenotypes from cross-modal embeddings of ECGs shown in Fig. 2c.

#### Increasing the number of unlabelled samples improves the prediction of MRI derived phenotypes from cross-modal ECG representations

We now analyze the relationship between the amount of labelled data for supervised learning, the amount of unlabelled data for cross-modal autoencoding, and the performance of supervised learning from cross-modal latent representations. Such an analysis is crucial for understanding the number of labelled and unlabelled data samples needed to build an effective cross-modal autoencoder for use in practice. In Fig. 2d, we focus on such an analysis for the practically relevant setting of predicting MRI derived phenotypes from ECGs. In particular, we measure the mean R^2^ performance across all 9 MRI derived phenotypes from Fig. 2c as a function of the number of unlabelled samples for autoencoding and labelled samples for supervised learning from cross-modal embeddings. Performing a scaling law analysis (see Methods 1.2), we observe that collecting unlabelled samples for autoencoding leads to roughly twice the increase in predictive performance as collecting labelled samples for supervised learning. Since the collection of unlabelled ECG-MRI pairs is easier than the collection of labelled MRIs, our cross-modal autoencoder is able to leverage easily collectable data to improve the performance on these downstream phenotype prediction tasks.

### Cross-modal Autoencoder Framework Enables Generating Cardiac MRIs from ECGs

Our framework enables the translation of ECGs, an easy-to-acquire modality, to cardiac MRIs, a more expensive, difficult-to-acquire modality. To perform such translation, we simply provide test ECGs into our ECG-specific encoder and then apply the MRI-specific decoder to translate from ECGs to MRIs. We note that since the two data modalities capture complementary cardiac features (ECGs capturing myoelectric information and MRIs capturing structural information), such translation is a nontrivial task. Nevertheless, we show that the translation of ECGs provided by a cross-modal autoencoder remarkably captures features present in MRIs, and we quantify the amount of such features captured via the translation.

#### Cardiac MRIs generated from test ECGs capture MRI specific phenotypes

We begin by qualitatively analyzing the reconstructions and translations of 12-lead ECG and 50 frame cardiac MRI test pairs using our cross-modal autoencoder. In Fig. 3, we demonstrate that translations from ECGs to MRIs generally capture MRI-derived phenotypes such as left ventricular mass (LVM) or right ventricular end-diastolic volume (RVEDV). In Fig. 3a and b, we consider translating from ECGs to MRIs for test samples of individuals with high or low LVM/RVEDV. We observe that the corresponding translations generally capture whether an individual has high or low LVM/RVEDV, as indicated by the annotated regions in red. For comparison, we additionally present reconstructions given by our model when provided the test MRI as an input. These reconstructions demonstrate that the MRI-specific decoder has the capacity to reconstruct fine grained details of an MRI. Hence, the difference in quality between reconstructions and translations can be attributed to the difference in embedding provided from ECG and MRI specific encoders. Additional translations from ECG to MRI (and vice-versa) are presented in Supplementary Figure S3. Supplementary Figure S4 demonstrates that decoding ECG or MRI cross-modal embeddings after shifting them in a direction of phenotypic effect (e.g. moving from low LVM to high LVM) leads to the desired phenotypic effect on the original modality (e.g. increased LVM in the corresponding generated MRI).

**Figure 3:**
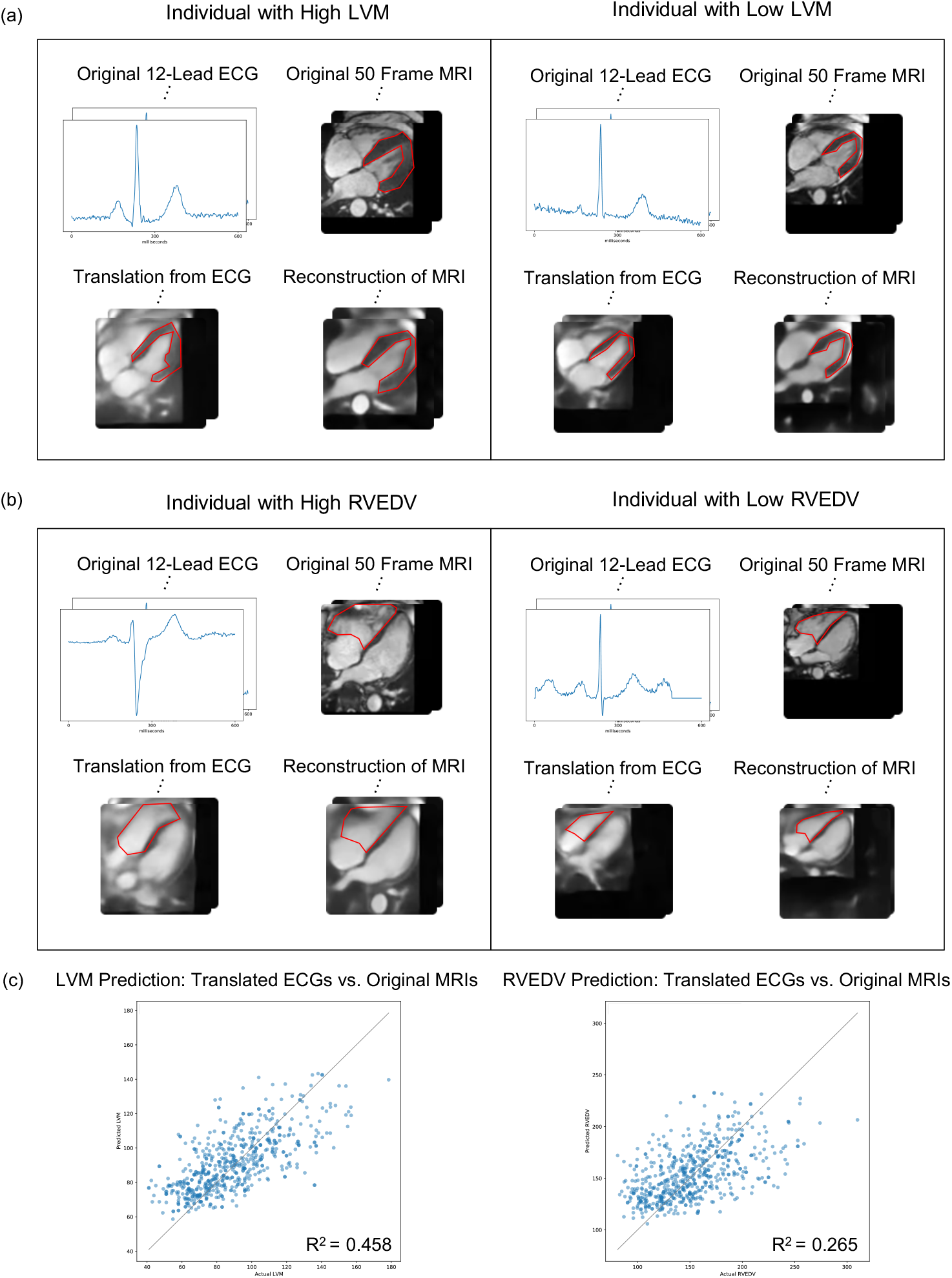
Cross-modal autoencoders enable imputing cardiac MRIs from ECGs while capturing MRI specific features such as left ventricular mass (LVM) and right ventricular end-diastolic volume (RVEDV) on test MRI-ECG pairs. (a) Examples showing qualitatively that MRIs imputed from test ECG samples capture LVM for those individuals with LVM in the highest and lowest quartile. The LVM in the original, translated, and reconstructed MRI is shown in red. (b) Examples showing qualitatively that MRIs imputed from test ECGs capture RVEDV for those individuals with RVEDV in the highest and lowest quartile. The RVEDV in the original, translated, and reconstructed MRI is shown in red. (c) The predictions of LVM and RVEDV on MRIs imputed from test ECGs correlate with the predictions of these phenotypes performed on the original MRIs.

In order to quantify the effectiveness of the translations using the cross-modal autoencoder, we compare the predictions of the translation to a neural network directly trained to predict LVM and RVEDV on the original modality. In particular, in Fig. 3c, we verify that the prediction of LVM and RVEDV from the reconstructed and translated MRIs positively correlates with that from ground truth MRIs. Hence, the translations of test ECGs provided by our cross-modal autoencoder indeed generally capture MRI derived phenotypes, as shown in Fig. 3a and b.

### Cross-modal Autoencoder Framework Enables Genome-wide Association Study using Integrated Latent Space

Next, we analyze whether cross-modal embeddings can be used to identify genotype-phenotype associations related to the heart. As a first step, we verify that performing a GWAS on labels derived from cross-modal representations leads to the recovery of SNPs previously associated with common disease phenotypes. We then develop a method based on the cross-model latent space to perform GWAS in the absence of labelled phenotypes, i.e. an unsupervised GWAS. We demonstrate that our unsupervised GWAS approach applied to cross-modal embeddings recovers SNPs typically identified by performing GWAS on labelled data, as well as those found in more computationally demanding ECG-wide screens [34].

#### GWAS of phenotypes predicted from cross-modal representations recovers phenotype-specific SNPs

In order to verify that cross-modal representations capture genetic associations with respect to a specific phenotype, we perform a GWAS on single trait predictions based on these representations; see Methods 1.4 for a description of performing such GWAS and a list of confounders considered. As an example, the Manhattan plot in Fig. 4a shows that such GWAS for body mass index (BMI) predicted from cross-modal embeddings identifies the gene FTO, which is known to have an effect on BMI and obesity risk [35, 36]. Similarly, performing a GWAS of right ventricular ejection fraction (RVEF) predicted from MRI cross-modal representations identifies lead SNPs corresponding to genes BAG3, HMGA2 and MLF1, which have all been previously associated with RVEF [37]. These results indicate that our learned representations are physiologically meaningful. Additional examples for ECG phenotypes derived from cross-modal embeddings are presented in Supplementary S5.

**Figure 4:**
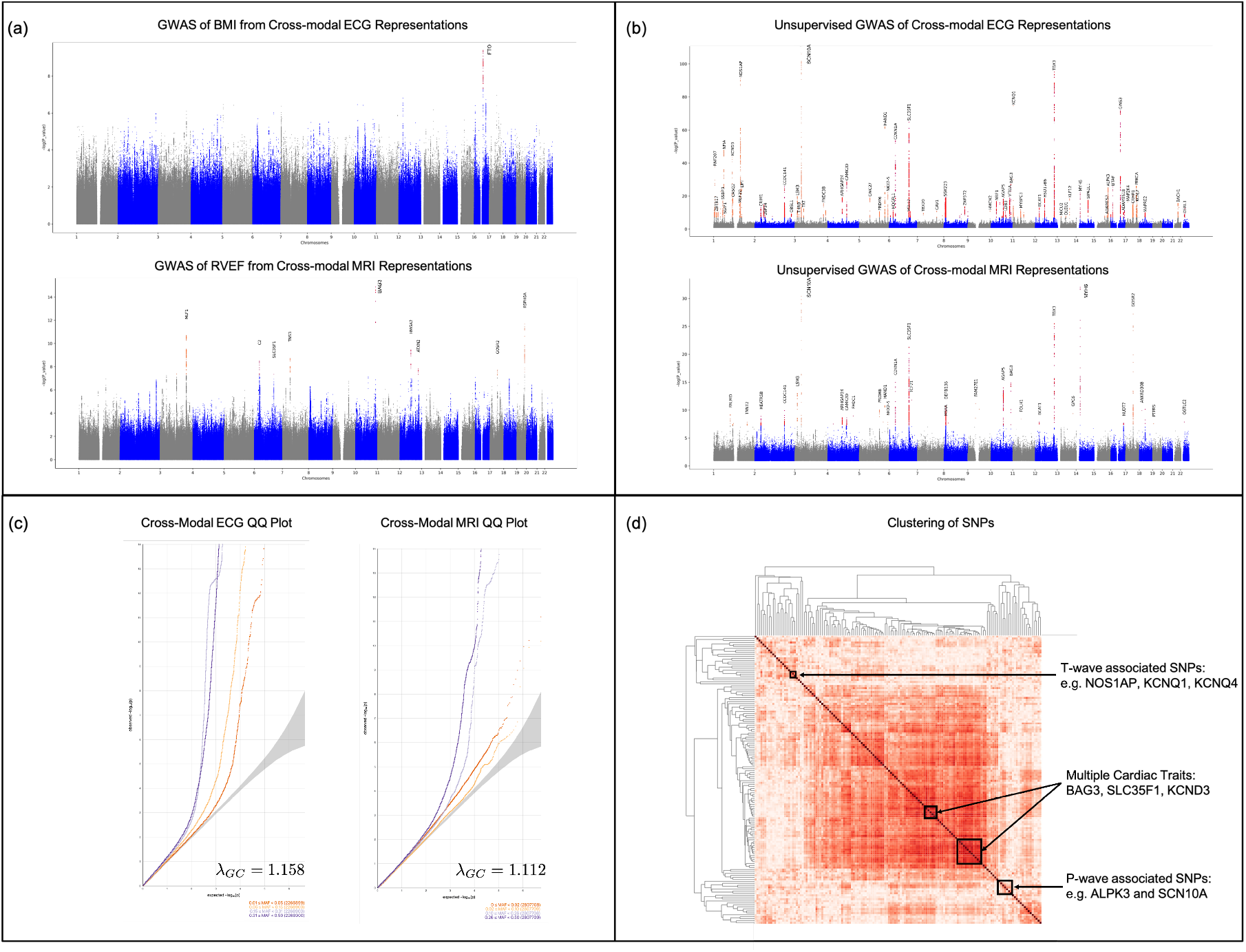
Cross-modal autoencoders capture genotype-phenotype associations for cardiovascular data. (a) Manhattan plots for GWAS of BMI and RVEF derived from cross-modal embeddings identify lead SNPs associated with these traits. For BMI, such GWAS identifies SNPs associated with FTO, which is known to have an effect on BMI and obesity risk. For RVEF, such GWAS identifies SNPs associated with BAG3, HMGA2, and MLF1, which have been previously associated with RVEF. (b) To more generally capture genetic associations with the heart, a GWAS can be performed in the cross-modal ECG and MRI latent space even in the absence of labelled data. The Manhatten plots of such unsupervised GWAS identify lead SNPs including those associated with NOS1AP, TTN, SCN10A, SLC35F1, KCNQ1, which have been previously associated with cardiovascular phenotypes. (c) The corresponding QQ plots and *λ*_*GC*_ factors indicate that there is minimal inflation in the unsupervised GWAS of cross-modal ECG and cardiac MRI embeddings. (d) Clustering SNPs by the direction from the mean embedding of homozygous reference samples to the mean embedding of heterozygous and homozygous alternate samples in order to group SNPs by similar phenotypic effect results in clusters of SNPs corresponding to those associated with the T-wave (NOS1AP and KCNQ1), the P-wave (SCN5A and ALPK3), as well as SNPs that affect multiple cardiac traits (e.g., BAG3, SLC35F1, and KCND3).

#### Unsupervised GWAS of cross-modal representations leads to the recovery of SNPs associated with a given modality

In order to characterize the genotype-phenotype associations from a modality more generally, we present a method for performing an “unsupervised GWAS” of cross-modal embeddings, i.e., a GWAS in the absence of any labelled phenotypes. Our approach is as follows: 1) for each SNP, we identify those individuals in the latent space that are either homozygous reference, heterozygous, or homozygous alternate; 2) we then ask whether these distributions are separable, and quantify the level of separation among these groups via a p-value from a Multivariate Analysis of Variance (MANOVA). The statistics used for computing p-values using MANOVA are discussed in Methods 1.4. Importantly, prior to performing MANOVA across the sets, we need to account for potential genetic confounders (stratification), which are inherently reflected in the cross-modal embeddings. To remove the effect of possible confounders, we use the iterated nullspace projection (INLP) method [38], which was used in natural language processing for removing features such as race or gender information from word embeddings. This method iteratively removes dimensions from the latent space until the remaining embeddings cannot be used to predict any confounder (see Methods 1.5 for additional details). After removing latent space dimensions that are predictive of confounders using INLP, we utilize MANOVA on the lower dimensional embeddings to perform an unsupervised GWAS (see Methods 1.5 for a list of confounders considered).

In Fig. 4b, we visualize the Manhattan plots resulting from utilizing our unsupervised GWAS approach on the cross-modal ECG and MRI embeddings. We observe that lead SNPs such as NOS1AP [39, 40], TTN [41, 42], SCN10A [43 –,45], SLC35F1 [46], and KCNQ1 [47, 48] are consistent with those identified by prior work. In Supplementary Fig. S6, we present the results of the unsupervised GWAS performed on the joint cross-modal embeddings of both ECG and MRI along with unsupervised GWAS performed on the unimodal autoencoder representations for these modalities. A full list of the lead SNPs identified in each analysis is presented in Supplementary Table S1. Furthermore, Fig. 4c shows the QQ plots and the corresponding λ_*GC*_ values to verify that the corresponding p-values are not inflated after removing the effect of confounders via INLP. In Supplementary Fig. S7, we analyze the impact of varying hyper-parameters of INLP on the level of inflation present in the resulting GWAS. As shown in Supplementary Fig. S8, the lead SNPs identified from our GWAS approach for cross-modal ECGs generally include those from GWAS on individual ECG phenotypes. We also identify many sites not previously associated with ECG or MRI traits, but which have clear associations with the cardiovascular system in general, for example NRP1, previously associated with HDL cholesterol [49], USP34 previously associated with cardiovascular disease [50], and NRG1 previously associated with systolic blood pressure [51]. We note that the cross-modal MRI GWAS identifies fewer lead SNPs than the cross modal ECG GWAS, which could be because MRIs are more strongly associated with the confounders and thus removing confounders via INLP may also remove genetic signal. Indeed, confounders such as age and sex are much more easily predicted from cross-modal MRI embeddings than cross-modal ECG, as is showcased in Fig. 2c. To illustrate the difference between between unsupervised GWAS of different representations, we compare the corresponding differences in Manhattan plots in Supplementary Fig. S9.

#### Clustering SNPs in the cross-modal latent space identifies SNPs with similar phenotypic impact

An additional benefit of our cross-modal approach for genetic discovery is that we can cluster SNPs in the latent space to group those with similar phenotypic effects. In particular, we perform hierarchical clustering based on the direction from the mean embedding of the homozygous reference group to the mean embedding of the heterozygous and homozygous alternate groups for any given SNP (see Fig. 1d). In Fig.4d, we analyze the SNP clusters given by performing hierarchical clustering on the SNP signatures in the cross modal embeddings given only ECG inputs (see Methods 1.4 for details regarding hierarchical clustering). In particular, we find two clusters corresponding to SNPs affecting the T-wave (SNPs associated with NOS1AP and KCNQ1) and P-wave (SNPs associated with SCN5A and ALPK3) of the ECG. We find several additional clusters corresponding to SNPs affecting multiple cardiac traits such as those associated with BAG3, SLC35F1, or KCND3. Supplementary Fig. S10 shows a high resolution version of this clustering and a clustering of a subset of lead SNPs, which illustrates robustness of our clusters.

## Discussion

In this work, we developed a cross-modal autoencoder framework for integrating data across multiple modalities to learn holistic representations of physiological state. Using the heart as a model system, we integrated cardiac MRI and ECG data to showcase the benefit of cross-modal representations via the following three applications: (1) improving prediction of phenotypes from a single modality; (2) enabling imputation of hard-to-acquire modalities like MRIs from easy-to-acquire ECGs; and (3) identifying genotype associations with general cardiovascular phenotypes. In particular, we showed that cross-modal representations improve prediction of cardiovascular phenotypes from ECGs alone. This setting is of practical importance given the abundance of ECG data over more difficult-to-acquire modalities such as MRI. Interestingly, we observed that increasing the number of unlabelled ECG and MRI pairs was more beneficial than increasing the number of labelled MRI data. We also demonstrated that cross-modal autoencoders enable imputing cardiac MRIs from ECGs. Importantly, we showed that the MRI-derived phenotypes are conserved in the translation. We also showed that the cross-modal representations can be used to perform GWAS. Notably, such an analysis not only recovers known phenotype-specific SNPs, but can also be used to perform unsupervised GWAS to identify SNPs that generally affect the cardiovascular system.

This novel framework for performing unsupervised GWAS in cross-modal representations opens important avenues for future work. Since cross-modal autoencoders learn representations from modalities directly, confounders are typically embedded in the representations. Indeed, we for example observed that MRI cross-modal embeddings can predict sex and age effectively. To minimize the effect of such confounders when performing genetic analyses, we were stringent in adjusting the latent space such that one could no longer predict confounders effectively from the learned representations. Developing more causally-grounded methods for confounder removal from a cross-modal latent space is an important open problem. Moreover, via a simple clustering of cross-modal embeddings, our framework allows for grouping SNPs by phenotypic effect without the need of labelled phenotypes. Since our framework can be used to integrate any number of data modalities, an exciting direction of future work is to use such modalities in other organs to better characterize the effect of SNPs with similar signatures in an unsupervised manner. Such identification requires reliable translation from the cross-modal latent space into the different modalities. While we showed that our framework is capable of translating from easy-to-collect ECGs to more difficult-to-collect MRIs while preserving relevant features, an interesting direction of future work is to understand how far such translations can be pushed.

With the rise of Biobanks around the world, our cross-modal integration framework opens an important novel avenue to integrate multiple modalities to build better representations of patient physiological state and thereby have an important impact on diagnostics and genomics. While we demonstrated the effectiveness of our cross-modal autoencoder framework on the cardiovascular system, our framework is broadly applicable to other organ systems.

## Methods

### 1.1 Study Design

All analyses were performed on the UK Biobank, a prospective cohort of over 500,000 healthy adults that were aged 40-69 during enrollment, which took place from 2006-2010. At the time of our analysis the UK Biobank had released cardiovascular magnetic resonance imaging for over 44,644 participants, 38,686 of whom also had a 12-lead 10-second resting ECG acquired on the same day. While different MRI views were obtained, we only considered the 4-chamber long axis view with balanced steady-state free-precession cines, containing 50 frames throughout the cardiac cycle. The ECG data also spanned a single cardiac cycle, because we used the 1.2-second 600-voltage median waveforms derived from the full 10-second ECG. All voltages were transformed to millivolts, and all MRI values were normalized to have mean 0 and standard deviation 1 for each individual. The MRIs were cropped to the smallest bounding box which contained all cardiac tissues in all 50 frames as determined by the semantic segmentation in [52].

### 1.2 Cross-modal Autoencoder Architecture and Training Details

#### Model architecture

The modality-specific encoders and decoders used in this work were selected through Bayesian hyper-parameter optimization [53]. In particular, we used a base architecture of densely connected parallel convolutional blocks [54, 55] with 1d convolutional layers for ECGs and 2d convolutional layers for MRIs. For modality-specific modals, we optimized over the width, depth, activation functions, regularization and normalization strategies to achieve minimum reconstruction error for a given maximum overall capacity of 10 million parameters and a 256 dimensional latent space. Since optimization occurs for each modality independently, encoding, decoding and pairing are distinct tasks and can be trained asynchronously and distributed across machines. We note that simpler architectures such as those from [56] are also usable in our framework, but we observed that the optimized models showed improvements in convergence speed, reconstruction, and latent space utility for downstream tasks (see Supplementary Fig. S1).

To ensure that only one modality is needed at test time, we additionally utilized dropout [57] to merge modality specific embeddings. In particular, during training, we employed dropout of a random subset of coordinates of the ECG embedding and merged it with the complementary coordinates from the MRI embedding. The resulting merged embedding was then decoded to reconstruct the original ECG and MRI examples.

#### Training methodology

Let 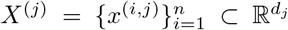 denote the set of samples of modality *j* for *j* ∈ [*m*] where [*m*] = {1, 2, … *m*}. Consider the paired setting where the samples 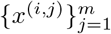 correspond to multiple data modalities for the same sample (e.g. cardiac MRI and ECG for the same patient). Given a subset of these modalities {*x*^(*i,j*)^}_*j*∈ ℐ_ for ℐ ⊂ [*m*], we constructed a cross-modal autoencoder that produces the remaining representations {*x*^(*i,j*)^}_*j*∈[*m*]− ℐ_ as follows. We decomposed our model into encoders 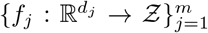 and decoders 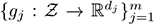, where the functions *f*_*j*_ and *g*_*j*_ are parameterized using deep neural networks. The neural networks were trained to both pair and reconstruct each data modality. Modality specific encoders and decoders allowed for inferring all modalities given any single one.

The training loss, ℒ for cross-modal autoencoders is given as the linear combination of the following two losses: (1) a reconstruction loss, *L*_*Rec*_, which is used to reconstruct the original modalities; and (2) a representation loss *L*_*Contrastive*_, which is used to ensure that the representations for modalities corresponding to the same sample are embedded nearby in the latent space. Formally, we have:

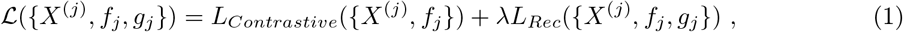

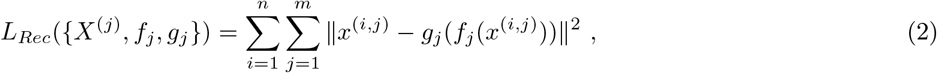

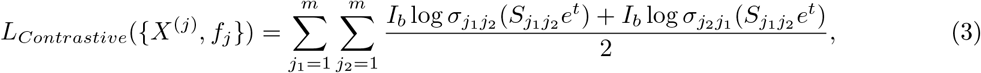

where 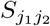 is the matrix of cross modality embedding similarities within each batch, given by

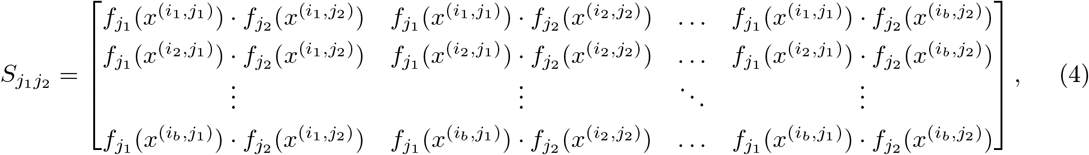

*σ* is the softmax function 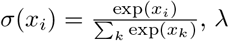 is a hyperparameter to balance the losses, *t* is a trainable temperature scalar as in [58], and *I*_*b*_ is the *b* × *b* identity matrix. We let 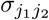 denote the softmax across rows 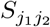 and 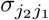 the columns of 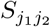. Intuitively, the contrastive loss above pushes embeddings from the same individual and different modalities closer together while pulling apart embeddings of different individuals and different modalities.

In our experiments, we used a batch size of 4 samples (*b* = 4) and used λ = 0.1. All models were optimized with the Adam optimizer [59] and a learning rate of 1e-3 for unimodal autoencoder training and 2e-5 for cross-modal fine-tuning. The learning rate was decayed by a factor of 2 after each epoch without an improvement in validation loss and after 3 decays optimization was terminated.

### 1.3 Models, Data, and Scaling Law for Phenotype Prediction Tasks

#### Supervised learning models for phenotype prediction tasks

We compared phenotype prediction from cross-modal embeddings to training supervised models with the same encoder architecture as described in Section 1.2. In particular, we trained these supervised models for phenotype prediction by adding a last layer and updating the weights via a logcosh loss for continuous tasks and cross entropy loss for categorical tasks. We also used the same optimization procedures for the hyper-parameters and the same stopping criteria as described in Section 1.2.

#### Data splits for phenotype prediction tasks

For all phenotype prediction tasks, we only considered data that was held out during cross-modal autoencoder training. This is crucial since otherwise the autoencoder would automatically utilize both MRI and ECG data for all phenotype predictions and thus naturally perform better than prediction from any individual modality. Since we were limited by the availability of labelled data for MRI derived phenotypes, we held out all data for which there was an available MRI derived phenotype from the autoencoder training and validation set. This left us with 4218 samples containing MRI derived phenotypes. For MRI derived phenotype prediction, we split these into 3163 samples for training, 527 for validation, and 528 for test. Only 4120 of these samples had corresponding ECG derived phenotypes, and so we used 3083 of these for training, 516 for validation, and 521 for test. For categorical general phenotypes, we used the same splits as those for MRI derived phenotypes. For continuous valued general phenotypes, we considered only the subset of the 4218 samples that had labels available. In particular, we used 3158 samples for training, 527 for validation, and 527 for testing.

#### Linear, logistic, and kernel regression models for phenotype prediction tasks

For phenotype prediction from latent space embeddings, we considered the performance of three models (1) kernel regression with the Neural Tangent Kernel (NTK) [60] ; (2) linear regression ; and (3) logistic regression. We considered the NTK since it was shown to have superior performance on supervised learning problems [61, 62]. For the prediction of MRI derived phenotypes, ECG derived phenotypes, or continuous general phenotypes, we measured performance using *R*^2^, and we compared the performance of the NTK and linear regression. For the prediction of categorical phenotypes, we measured performance using the area under the receiver operator characteristic curve (AUROC), and we compared the performance of the NTK and logistic regression. We utilized EigenPro [63] to solve kernel ridge-less regression and linear regression. We used the validation splits to select the early-stopping point for the EigenPro iteration. Similar results can be obtained with 𝓁_2_-regularized kernel and linear regression using the Scikit-learn implementation [64], but require the more computationally demanding step of fine-tuning of the regularization parameter based on validation performance. For classification tasks, we used the implementation of 𝓁_2_-regularized logistic regression from [64], and we applied the following weighting on the loss to account for class imbalances: if there were *n* total samples of which *r* had label 1, then we weighted the loss for these samples by 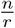 and the loss for the samples with label 0 by 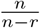..

#### Scaling law for prediction of MRI-derived phenotypes from cross-modal ECG representations

We now describe our scaling law analysis used to determine the relationship between the amount of labelled data for supervised learning (denoted by *v*), the amount of unlabelled data for crossmodal autoencoding (denoted by *u*), and the performance of supervised learning from cross-modal latent representations (denoted by *r*). We used linear regression to map from (log_2_ *v*, log_2_ *u*) to *r* for the 54 samples considered in Fig. 2d. The corresponding linear mapping is given by:

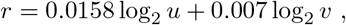

and yields *R*^2^ = .983. Hence for these tasks, we were able to reliably predict the boost in performance from supervised models on cross-modal embeddings when varying the number of unlabelled ECG-MRI pairs and labelled MRIs. Note that the coefficient of log_2_ *u* is over twice that of log_2_ *v* implying that collecting unlabelled ECG-MRI pairs leads to roughly twice the increase in predictive performance as collecting labelled samples.

### 1.4 GWAS of Phenotypes Derived from Cross-modal Representations

#### GWAS on phenotypes predicted from latent representations

We found multi-pathway genetic signals in the cross modal latent spaces by analyzing the inferences of the kernel regression models described above. Specifically, we trained ridge-regression models to use modality-specific cross modal embeddings to predict ECG phenotypes (eg PQ Interval N=36,645), MRI-derived phenotypes (e.g. RVEF N=4788) and general demographics (e.g. BMI N=38,000). These simple models endow these GWAS with much greater statistical power, since phenotypes can be predicted for the whole cohort, not just those with labels, as described in [65]. For example, GWAS of the less than 5,000 MRI-phenotypes returned by [33] yield no genome wide significant hits, while inferences from ridge-regression yield dozens of plausible sites. These sites are confirmed by GWAS of traits computed from semantic segmentation described in [37]. Models were fit with 80% of the available labels and and evaluated on the remaining 20% and then inferred on the entire cohort.

#### Confounders considered in GWAS

To account for population structure and ascertainment biases all GWAS were adjusted for the top 20 principal components of ancestry, the UK Biobank assessment center where the measurements were conducted, the genomic array batch, as well as age and sex of each individual.

### 1.5 Unsupervised GWAS of Cross-modal Representations

#### Application of iterative nullspace projection for removing confounders

To remove the effect of confounders, we utilized the idea of iterative nullspace projection from [38]. Intuitively, this algorithm reduces the dimensionality of the latent space by removing dimensions that are useful for the prediction of confounders. Unlike the original implementation, which is designed primarily for categorical confounder removal and has additional memory overhead from storing projection matrices, we here present an implementation for continuous confounder removal that avoids extra overhead by utilizing the singular value decomposition (SVD). At a high-level, the algorithm involves iterating the following steps until the *R*^2^ from step 1 is below a pre-selected threshold (we used *R*^2^ < 0.001).

*Step 1:* Use linear regression to learn a mapping from cross-modal latent embeddings to confounders.

*Step 2:* Use the singular value decomposition to construct a projection matrix that projects onto the directions of the cross-modal space that are least useful for confounder prediction, i.e. the nullspace of the predictor from step 1.

*Step 3:* Multiply the cross-modal embeddings by the projection matrix found in step 2.

Mathematically, these steps are implemented as follows. Let 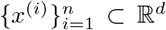 denote the cross-modal latent space embeddings for *n* individuals and let 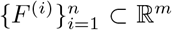 denote the set of *m* confounders for the *n* individuals. To correct for confounders, we do the following:

*Step 1:* Learn the regression coefficients *w* ∈ ℝ^*m*×*d*^ by minimizing the loss:

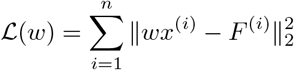

*Step 2:* Let *w* = *U* Σ*V* ^*T*^ given by the SVD, where *V* ^*T*^ ∈ ℝ ^*d*×*d*^. To project out the components corresponding to the confounders, select out the bottom *d* − *m* rows of *V* ^*T*^ into a matrix 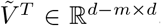.

*Step 3:* Replace each *x*^(*i*)^ with 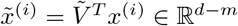.

Repeat the above steps until the *R*^2^ of the predictor *w* is lower than a fixed threshold.

#### MANOVA p-value computation

The p-values reported for unsupervised GWAS are from Pillai’s trace test statistic from the MANOVA computation. The Python statsmodels [66] package was used to perform MANOVA.

#### Clustering of SNPs by effect

Agglomerative clustering with Ward’s method, which minimizes the total within-cluster variance, was applied to the matrix of SNP vectors. The python sklearn [64] clustering package was used to derive the clusters and dendrograms.

## Data Availability

UK Biobank data is available to researchers from research institutions with genuine research inquiries, following IRB and UK Biobank application approval. All GWAS summary statistics will be available upon publication and are peruse-able on LocusZoom.

## Code Availability

Serialized encoders, decoders and full autoencoder models will be made available in the ML4H Model Zoo in the github repository: https://github.com/broadinstitute/ml4h [67].

