## Supplementary Materials for "A Cross-Modal Autoencoder Framework Learns Holistic Representations of Cardiovascular State"

**This PDF file includes:**

Supplementary Figures S1 to S10
Supplementary Table S1
Supplementary Videos S1 to S2

### Supplementary Figures

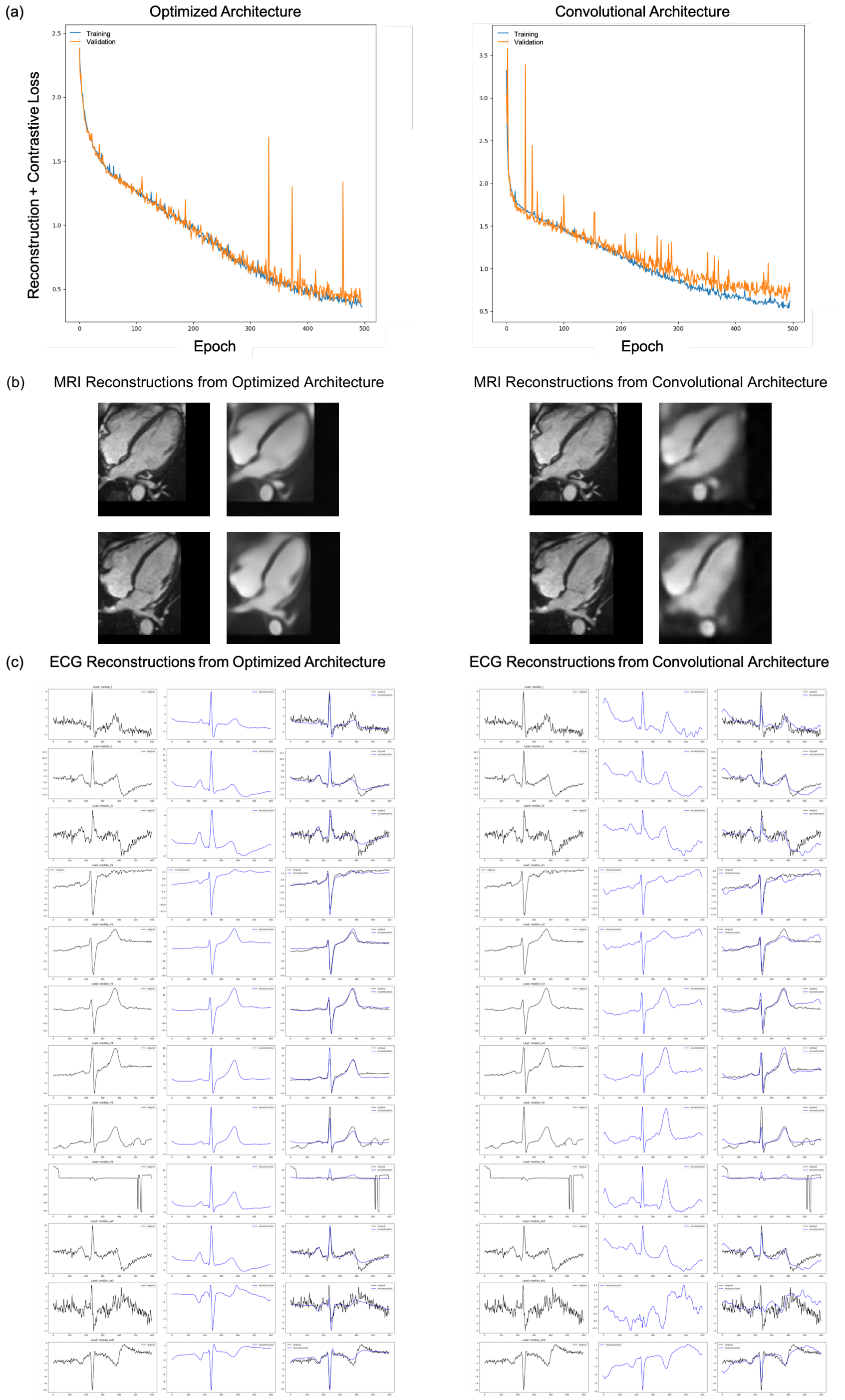


Supplementary Fig. S1: A comparison of our hyper-parameter optimized architecture with a standard convolutional architecture from [1]. (a) We visualize the training and validation loss across 500 epochs and observe that the hyper-parameter optimized architecture produces lower loss than the CNN Autoencoder architecture. (b-c) We visualize the reconstructions of test MRI and ECG samples from the optimized architecture and convolutional architecture and observe that reconstructions from the optimized architecture are higher quality (lower loss) than those from the architecture from [1].

(a) $R^2$ Values for ECG-Derived Phenotype Prediction from Cross-modal MRI Embeddings

| Model\Phenotype | PQ Interval | QT Interval | QTC Interval | QRS Duration | RR Interval | Average |
|---|---|---|---|---|---|---|
| Kernel Regression | 0.51 | 0.26 | 0.19 | 0.25 | 0.49 | 0.34 |
| Linear Regression | 0.50 | 0.25 | 0.18 | 0.24 | 0.49 | 0.33 |

$R^2$ Values for MRI-Derived Phenotype Prediction from Cross-modal ECG Embeddings

| Model\Phenotype | LVM | LVEDV | LVEF | LVESV | LVSV | RVEF | RVESV | RVSV | RVEDV | Average |
|---|---|---|---|---|---|---|---|---|---|---|
| Kernel Regression | 0.53 | 0.45 | 0.11 | 0.39 | 0.31 | 0.13 | 0.44 | 0.31 | 0.48 | 0.35 |
| Linear Regression | 0.51 | 0.43 | 0.11 | 0.36 | 0.30 | 0.14 | 0.43 | 0.30 | 0.47 | 0.34 |

(b) $R^2$ Values for General Numerical Phenotype Prediction from Cross-modal ECG Embeddings

| Model\Phenotype | BMI | Age | Average |
|---|---|---|---|
| Kernel Regression | 0.36 | 0.27 | 0.32 |
| Linear Regression | 0.35 | 0.24 | 0.29 |

$R^2$ Values for General Numerical Phenotype Prediction from Cross-modal MRI Embeddings

| Model\Phenotype | BMI | Age | Average |
|---|---|---|---|
| Kernel Regression | 0.48 | 0.42 | 0.45 |
| Linear Regression | 0.47 | 0.40 | 0.44 |

(c) AUROC Values for General Categorical Phenotype Prediction from Cross-modal ECG Embeddings

| Model\Phenotype | Sex | Hypercholesterolemia | Hypertension | Average |
|---|---|---|---|---|
| Kernel Regression | 0.96 | 0.64 | 0.69 | 0.76 |
| Logistic Regression | 0.90 | 0.56 | 0.63 | 0.70 |

AUROC Values for General Categorical Phenotype Prediction from Cross-modal MRI Embeddings

| Model\Phenotype | Sex | Hypercholesterolemia | Hypertension | Average |
|---|---|---|---|---|
| Kernel Regression | 0.99 | 0.66 | 0.75 | 0.80 |
| Logistic Regression | 0.95 | 0.58 | 0.66 | 0.73 |

Higher $R^2$ and AUROC are better with a maximum value of 1.

Supplementary Fig. S2: A comparison of kernel, linear, and logistic regression models used for phenotype prediction from cross-modal ECG and MRI embeddings. Overall, we observe that kernel regression models outperform linear and logistic regression models for the tasks considered in Fig.2 of the main text. (a, b) We report $R^2$ for kernel and linear regression used in prediction of continuous valued phenotypes considered in Fig.2 of the main text. (c) We report AUROC for kernel and logistic regression used in prediction of categorical phenotypes considered in Fig.2 of the main text.

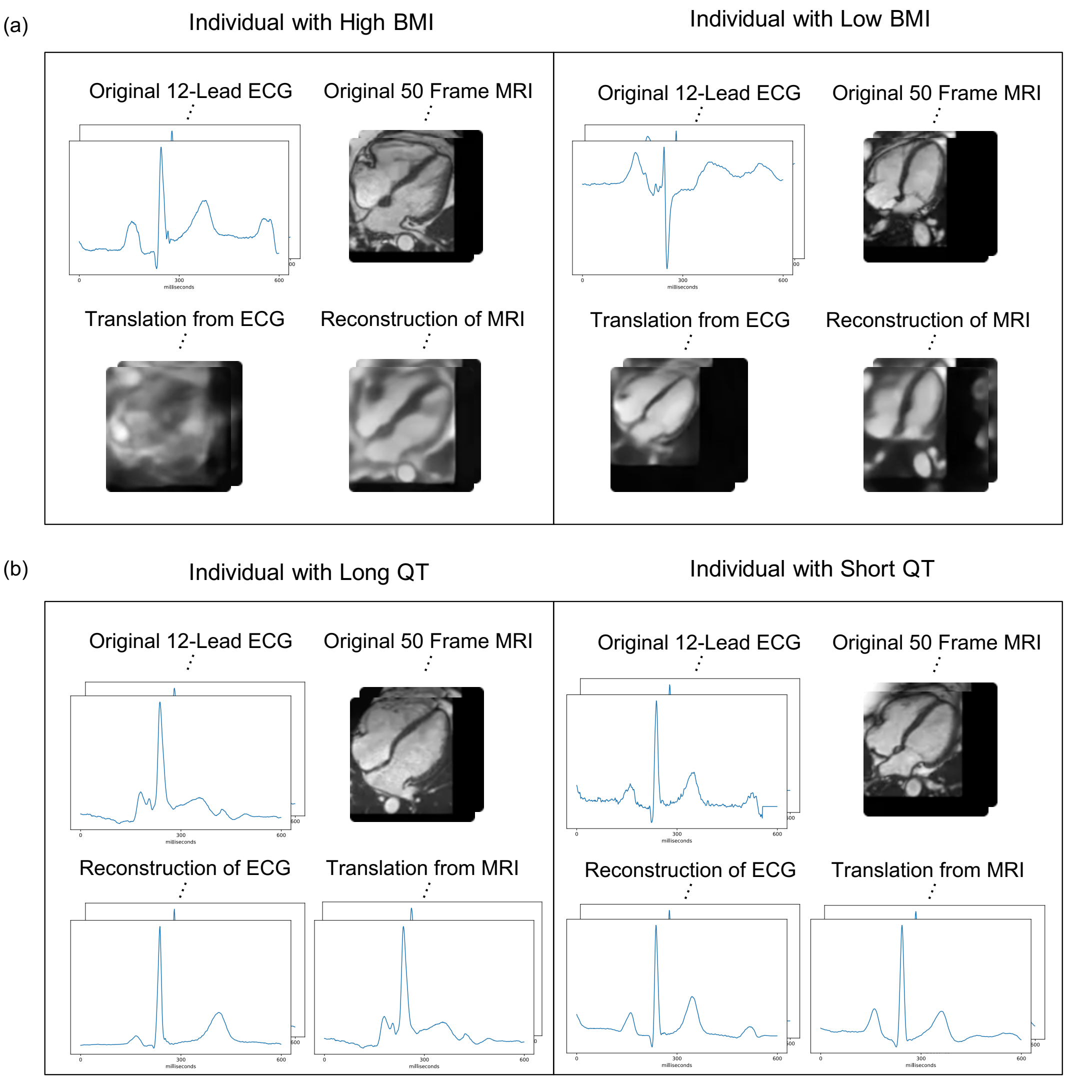


Supplementary Fig. S3: Additional examples of modality translation using cross-modal autoencoders. (a) Translation of ECG to MRI for individuals with high and low BMI. (b) Translation of MRI to ECG for individuals with long and short QT intervals.

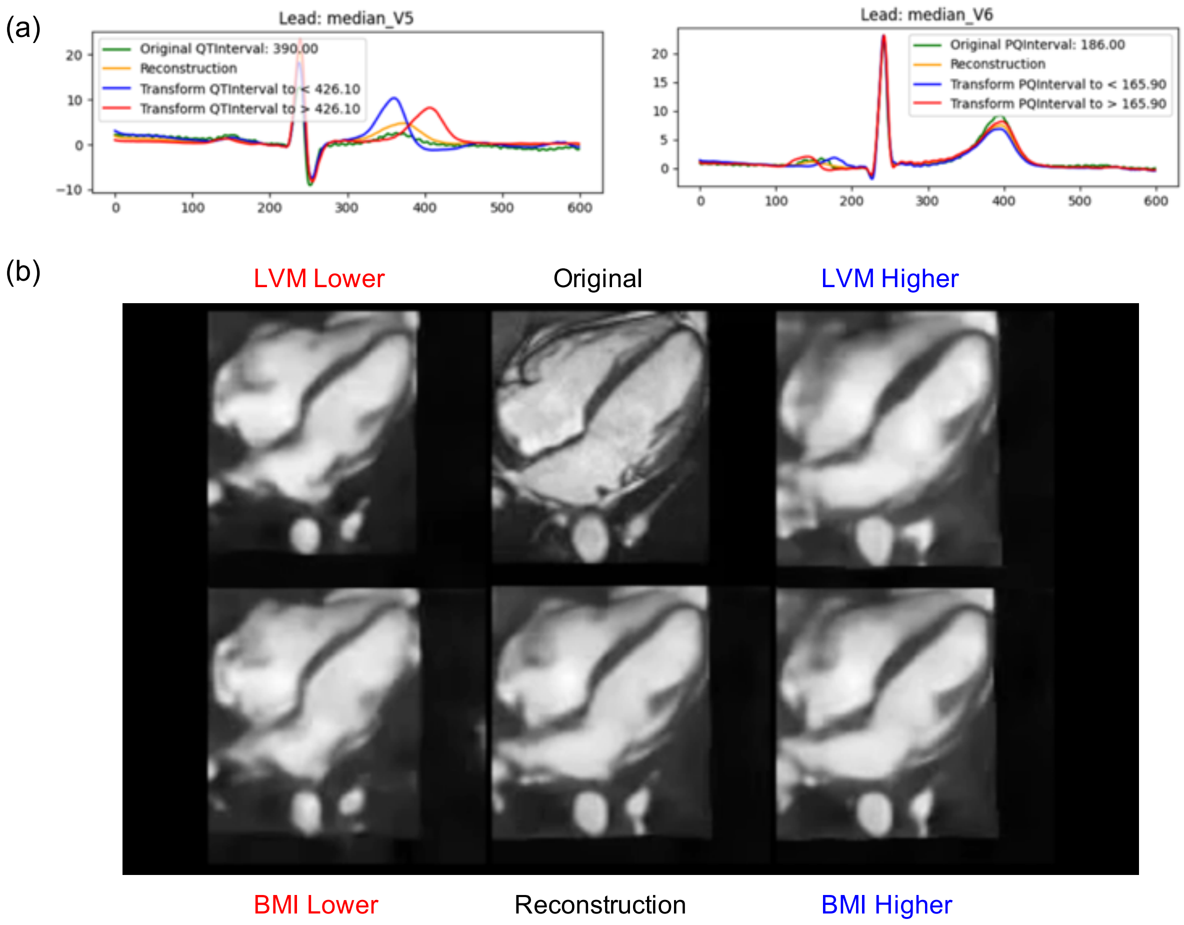


Supplementary Fig. S4: Translating cross-modal embeddings along a phenotype direction produces phenotype-specific impacts on ECGs and MRIs after decoding. (a) Translating cross-modal embeddings along the direction from short to long (or long to short) QT or PQ interval leads to corresponding increases (or decreases) of these intervals on the original ECGs. (b) Translating cross-modal embeddings along the direction from low to high (or high to low) LVM or BMI leads to corresponding increases (or decreases) of these phenotypes on the original MRI.

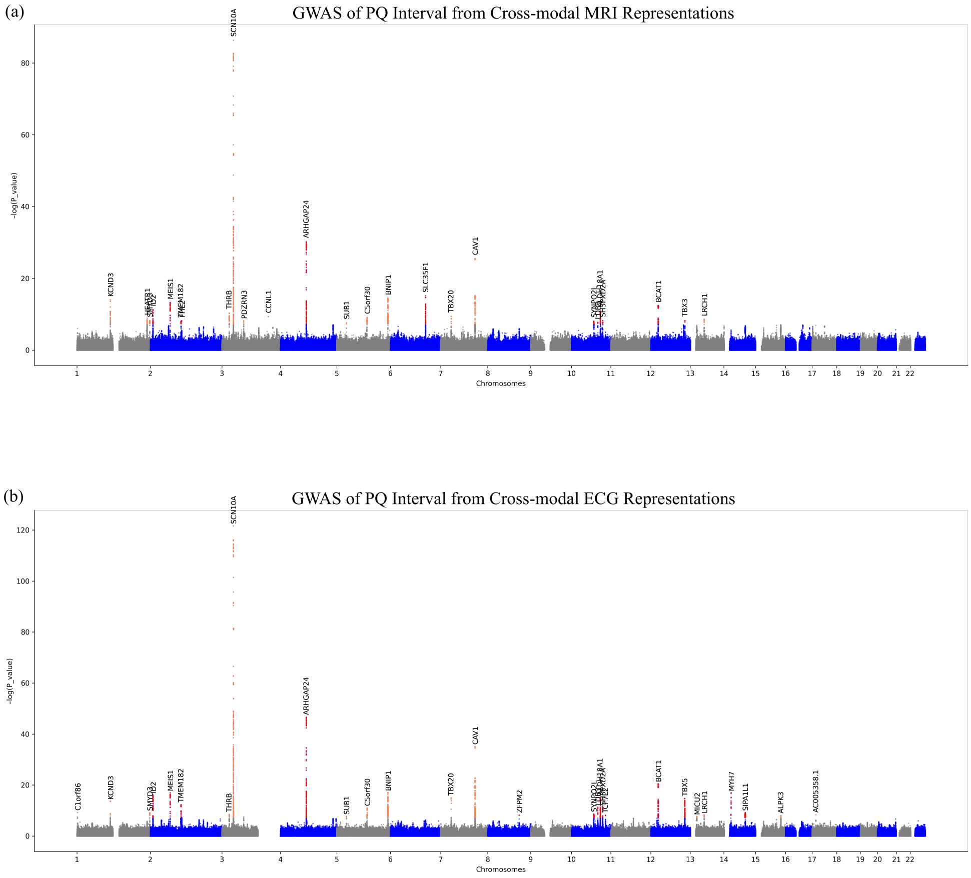


Supplementary Fig. S5: GWAS of PQ interval predicted from (a) MRI cross-modal representations or (b) ECG cross-modal representations identifies genes associated with PQ interval duration, including SCN10A, KCND3, and CAV1.

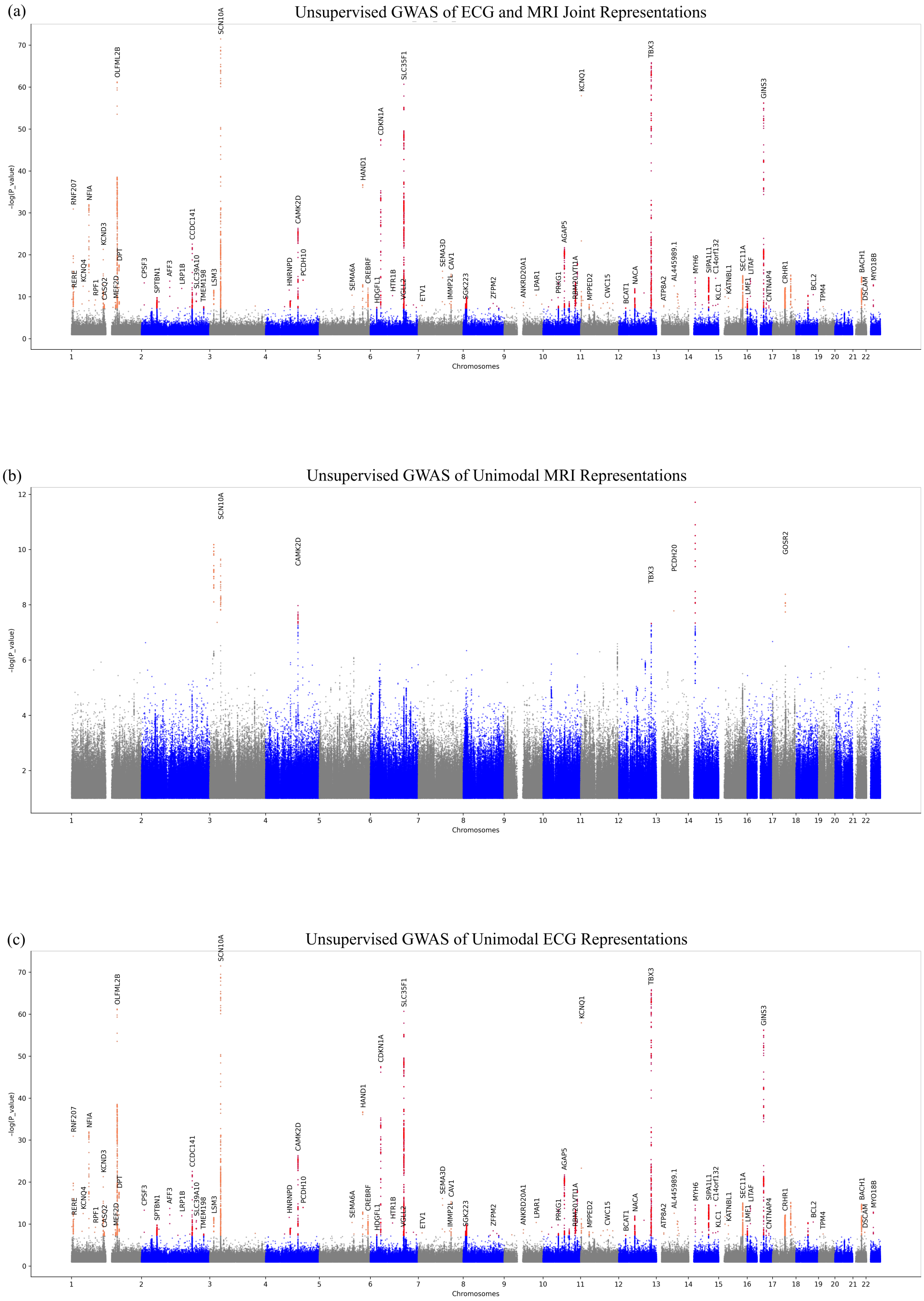


Supplementary Fig. S6: Unsupervised GWAS of cross-modal representations and representations from unimodal autoencoders. (a) Unsupervised GWAS of cross-modal latent representations of both ECG and MRI, i.e. representations corresponding to the instances where both ECG and MRI pairs are available as input to the model. (b) Unsupervised GWAS of MRI representations from a unimodal autoencoder. (c) Unsupervised GWAS of ECG representations from a unimodal autoencoder.

| Model | Number of Principal Components of Ancestry | INLP $R^2$ Threshold | Remaining Latent Dimensionality | Number of Lead SNPs Identified | GC $\lambda$ |
|---|---|---|---|---|---|
| Cross-modal ECG | 30 | 0.001 | 38 | 48 | 1.083 |
| Cross-modal ECG | 10 | 0.002 | 111 | 91 | 1.172 |
| Cross-modal ECG | 5 | 0.002 | 131 | 723 | 1.333 |
| Cross-modal ECG | 5 | 0.01 | 165 | 2720 | 2.72 |
| Unimodal ECG | 40 | 0.001 | 13 | 50 | 1.151 |
| Unimodal ECG | 30 | 0.001 | 49 | 97 | 1.228 |
| Unimodal ECG | 20 | 0.001 | 85 | 304 | 1.338 |
| Cross-modal MRI (256 latent dims.) | 10 | 0.002 | 136 | 26 | 1.086 |
| Cross-modal MRI (512 latent dims.) | 30 | 0.001 | 202 | 73 | 0.984 |
| Unimodal MRI | 10 | 0.002 | 167 | 6 | 1.0 |

(a)

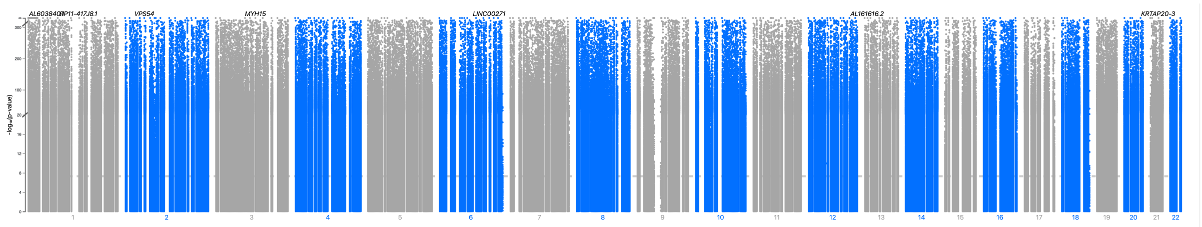

(b)

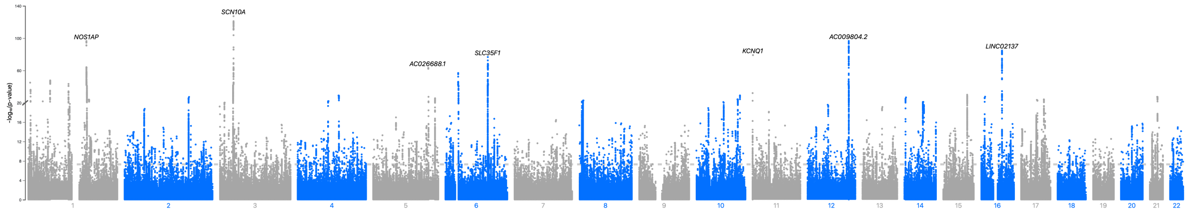


(c)

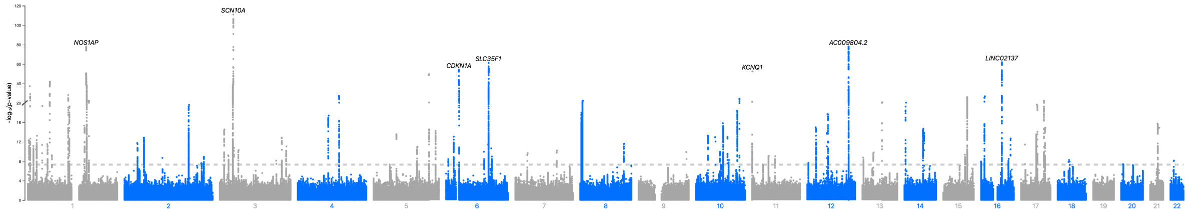


(d)

Supplementary Fig. S7: Impact of varying the number of principal components of ancestry and the threshold for iterative nullspace projection (INLP) on the number of lead SNPs recovered by unsupervised GWAS for cross-modal and unimodal ECG embeddings. (a) Using too few principal components (PCs) of ancestry or using too large of an $R^2$ threshold yield unsupervised GWAS that are inflated, as is indicated by that GC $\lambda$ values. (b) Manhattan plot for uncorrected GWAS, which is highly inflated. (c) Manhattan plot for GWAS corrected with 5 PCs and an INLP threshold of 0.01, which is again inflated. (d) Manhattan plot for corrected GWAS with 20 PCs and an INLP threshold of 0.0015, which is no longer inflated.

(a) Comparison of Lead SNPs Across Embeddings/Phenotypes

| | PQ Interval | QRS Duration | QT Interval | QTC Interval | RR Interval |
|---|---|---|---|---|---|
| Cross-modal ECG + MRI | 0.50 | .96 | 0.61 | 0.62 | 0.45 |
| Unimodal ECG | 0.55 | 0.91 | 0.67 | 0.62 | 0.45 |
| PQ Interval | 1.00 | 0.22 | 0.11 | 0.08 | 0.45 |
| QRS Duration | 0.23 | 1.00 | 0.11 | 0.17 | 0.18 |
| QT Interval | 0.09 | 0.09 | 1.00 | 0.5 | 0.45 |
| QTC Interval | 0.09 | 0.17 | 0.67 | 1.00 | 0.18 |
| RR Interval | 0.23 | 0.09 | 0.28 | 0.08 | 1.00 |

(b) Number of Lead SNPs from GWAS

| Cross-modal ECG + MRI | Unimodal ECG | PQ Interval | QRS Duration | QT Interval | QTC Interval | RR Interval |
|---|---|---|---|---|---|---|
| 93 | 86 | 22 | 23 | 18 | 24 | 11 |

Supplementary Fig. S8: (a) Unsupervised GWAS of cross-modal representations identifies several lead SNPs associated with the heart and includes those found from GWAS on ECG derived phenotypes. Entry $(i, j)$ of the table represents the ratio of the intersection of lead SNPs identified from GWAS using the representation in row $i$ and those identified from GWAS using the phenotype in column $j$ divided by the number of SNPs identified from GWAS using the phenotype in column $j$. We observe that lead SNPs identified by unsupervised GWAS of cross-modal ECG representations includes several of those from GWAS of PQ interval, QRS duration, QT interval, QTC interval, and RR interval. On the other hand, GWAS based of specific phenotypes (e.g. PQ interval, QRS duration, etc.) identifies lead SNPs that do not overlap much with those from GWAS of other ECG derived phenotypes. (b) A count of the number of lead SNPs identified by our unsupervised GWAS compared to GWAS on labelled ECG phenotypes. Our method recovers many more significant SNPs and includes those found via traditional GWAS approaches.

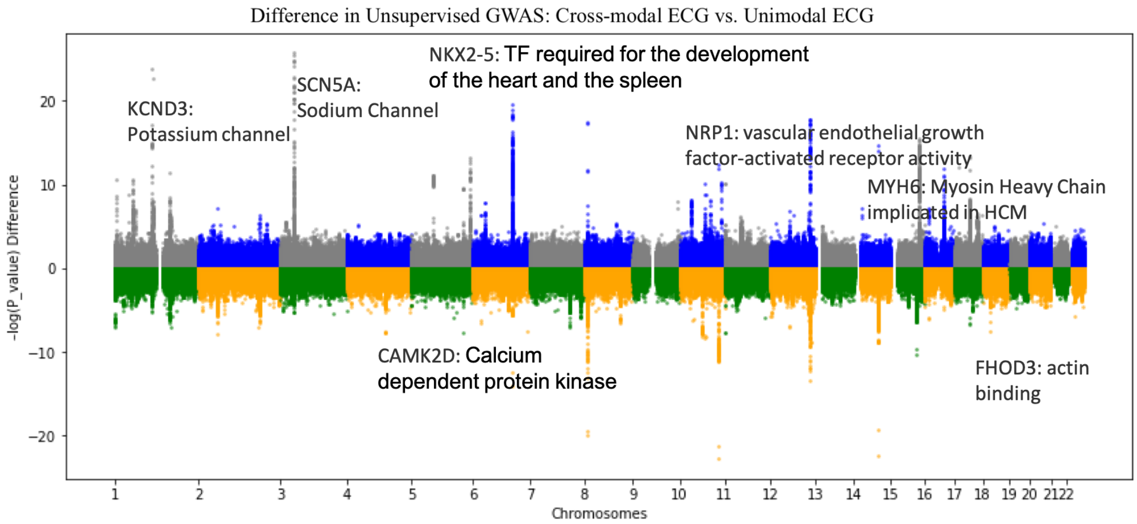


(a)

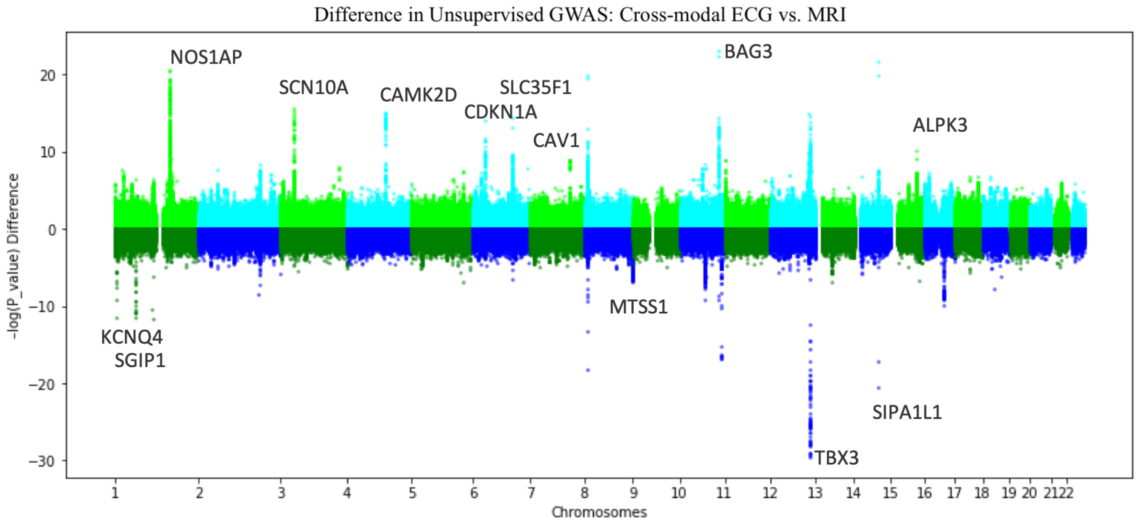


(b)

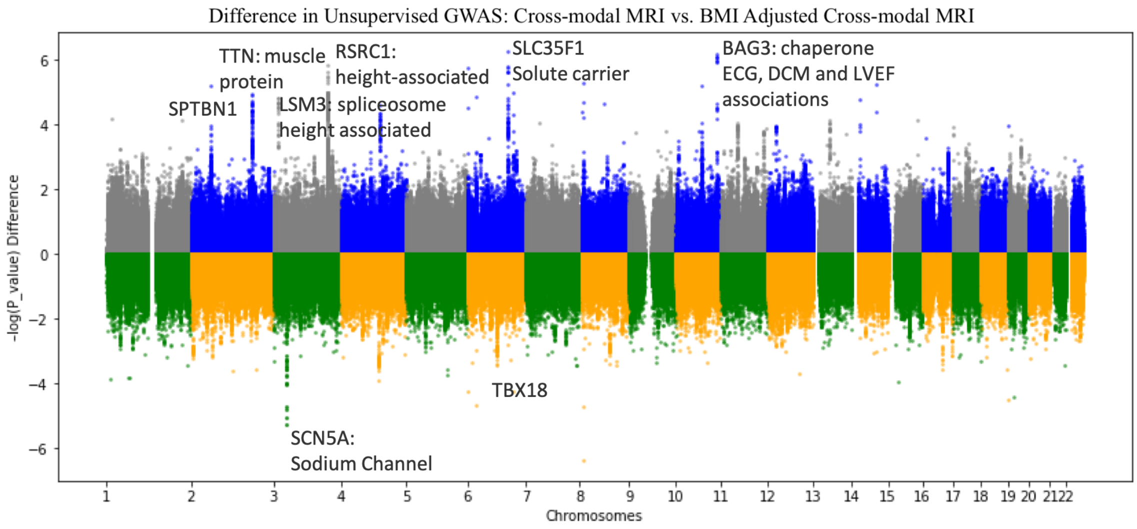


(c)

Supplementary Fig. S9: A visualization of the differences between Manhattan plots resulting from unsupervised GWAS. (a) The difference between unsupervised GWAS for cross-modal ECG representations vs. unimodal ECG representations. (b) The difference between unsupervised GWAS of cross-modal ECG representations and cross-modal MRI representations. (c) The difference between unsupervised GWAS for cross-modal MRI representations and BMI-adjusted cross-modal MRI representations.

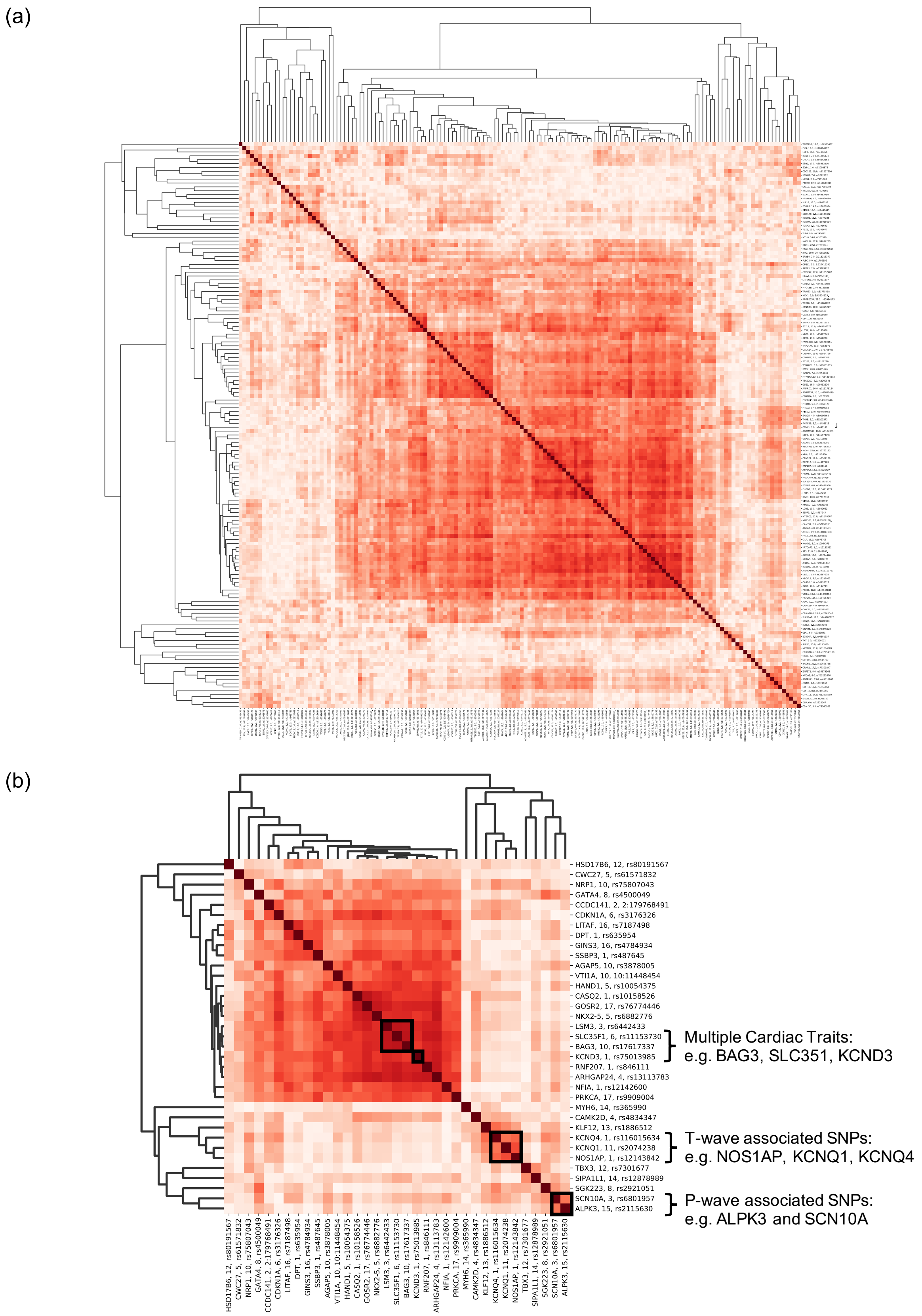


Supplementary Fig. S10: (a) Hierarchical clustering of SNPs by signature, i.e. the direction from the mean embedding of homozygous reference samples to the mean embedding of heterozygous and homozygous alternate samples. Darker colors indicate highly correlated SNP signatures. Several clusters arise including those corresponding to T-wave specific genes, P-wave specific genes, and genes with effects of multiple cardiac traits. (b) Hierarchical clustering on a smaller subset of lead SNPs identifies robustness of clusters by showing that the SNPs fall into the same phenotypic clusters as in (a).
